# Lineage and Spatial Mapping of Glioblastoma-associated Immunity

**DOI:** 10.1101/2020.06.01.121467

**Authors:** Vidhya M. Ravi, Nicolas Neidert, Paulina Will, Kevin Joseph, Julian P. Maier, Jan Kückelhaus, Lea Vollmer, Jonathan M Goeldner, Simon P. Behringer, Florian Scherer, Melanie Boerries, Marie Follo, Tobias Weiss, Daniel Delev, Julius Kernbach, Pamela Franco, Nils Schallner, Christine Dierks, Maria Stella Carro, Ulrich G. Hofmann, Christian Fung, Jürgen Beck, Roman Sankowski, Marco Prinz, Oliver Schnell, Dieter Henrik Heiland

## Abstract

The diversity of molecular states and cellular plasticity of immune cells in the glioblastoma environment is still poorly understood. Here, we performed scRNA sequencing of the immune compartment and mapped potential cellular interactions leading to an immunosuppressive microenvironment and dysfunction of T cells. Through inferring the dynamic adaptation during T cell activation, we identified three different terminal states with unique transcriptional programs. Modeling of driver genes for terminal T cell fate identified IL-10 signaling alterations in a subpopulation of HAVCR2(+) T cells. To explore in depth cellular interactions, we established an *in-silico* model by the integration of spatial transcriptomic and scRNA-sequencing, and identified a subset of HMOX1^+^ myeloid cells defined by IL10 release leading to T cell exhaustion. We found a spatial overlap between HMOX(+) myeloid and HAVCR2(+) T cells, suggesting that myeloid-lymphoid interaction causes immunosuppression present in tumor regions with enriched mesenchymal gene expression. Using human neocortical GBM model, coupled with patient-derived T cells, we confirmed that the functional interaction between myeloid and lymphoid cells, leads to a dysfunctional state of T cells. This IL-10 driven T cell exhaustion was found to be rescued by JAK/STAT inhibition. A comprehensive understanding of the cellular states and plasticity of lymphoid cells in GBM will aid towards successful immunotherapeutic approaches.

## Introduction

Tumor infiltrating lymphocytes, along with resident and migrated myeloid cells, account for a significant part of the tumor microenvironment in glioblastoma^1–3^. Most recently, the characterization of the myeloid cell population using scRNA-sequencing revealed remarkable heterogeneity with regards to cellular diversity and plasticity within the myeloid compartment^1,4^. However, the diversity of lymphoid cell types within malignant brain tumors remains unexplored and needs to be illuminated. Insights into the heterogeneity of cell type composition and driver genes for lineage differentiation within lymphoid compartment will aid in providing successful approaches for immunotherapy in the future. In other cancer entities such as colorectal cancer^5^, liver cancer^6^ or melanoma^7^, different T cell states have been investigated. Situations which involve prolonged immune activation and ambiguous stimulation, such as uncontrolled tumor growth or chronic infections, impede the ability of CD8^+^ lymphocytes to secrete proinflammatory cytokines and maintain their cytotoxic profile^7–9^. This cellular state, named dysfunctional or "exhausted" CD8^+^ lymphocytes, represents a paramount barrier to successful immune-vaccination or checkpoint therapy^2,10,11^. T cell exhaustion is partially orchestrated by regulation of inhibitory cell surface receptors (PD-1, CTLA-4, LAG-3, TIM-3 and others), in addition to anti-inflammatory cytokines such as IL-10 and TGF-ß. Glioblastoma, a common and very aggressive primary brain tumor in adults, is archetypical for tumors with a strong immunosuppressive microenviroment^12^. Current, immunotherapeutic approaches such as PDL1/PD1 checkpoint blockade^13^ or peptide vaccination^14^, led to remarkable responses in several cancers, has failed to demonstrate its effectivity in patients suffering from glioblastoma. To address the sparse knowledge with respect to the lymphoid cell population in glioblastoma, we performed deep transcriptional profiling by means of scRNA-sequencing, and mapped potential cellular interactions and cytokine responses that could lead to the dysfunctional and exhausted phenotype of T cells. Pseudotime analysis revealed an increased response to Interleukin 10 (IL10) during the transformation of T cells from the effector state to the dysfunctional state. To computationally explore “connected” cells driving this transformation, we introduced a novel approach termed “*nearest functionally connected neighbor* (NFCN)”, which identified a subset of myeloid cells marked by *CD163^+^* and *HMOX1^+^* expression. Furthermore, we performed spatially resolved transcriptomics, which confirmed the spatial overlap of exhausted T cells with *HMOX1^+^* myeloid cells, within regions of the tumor enriched with mesenchymal transcriptional signatures. Furthermore, using human neocortical GBM model with/without myeloid cell depletion, along with autografted T cell stimulation, we were able to conclusively validate our findings from the computational approach, which confirmed the role of myeloid cells as a key driver of the immunosuppressive microenvironment.

## Results

### Single cell Analysis of the Immune Cell Compartment in Glioblastoma

In order to interrogate the diversity of the immune microenvironment in glioblastoma, we performed droplet based 10X single cell sequencing of tissue samples from 8 patients, diagnosed with Glioblastoma. Lymphoid and myeloid populations (CD45^+^/CD3^+^) were sorted from neoplastic tissue specimens (**Figure 1a and Supplementary Figure 1a)**. The scRNA-seq data consisted of 47,284 cells, with a median number of 2,301 unique molecular identifiers (UMIs) and approximately 1023 uniquely expressed genes per cell. We corrected the data for mitochondrial genes, regressed out cell cycle effects and removed batch effects due to technical artifacts. We then decomposed the eigenvalue frequencies of the first 100 principal components and determined the number of non-trivial components by comparing them to randomized expression values, resulting in 41 meaningful components. Shared nearest neighbor (SNN) graph clustering resulted in 21 clusters (C0-C20) containing uniquely expressed genes. The major observed cell type when using the semi-supervised subtyping algorithm of scRNA-seq (SCINA-Model)^15^ and SingleR^16^ are: microglia cells (*TMEM119, CX3CR1* and *P2RY12*) and macrophages (*AIF1, CD68, CD163* and low expression of *TMEM119, CX3CR1*), followed by CD8^+^ T cells (*CD8A, CD3D*), natural killer cells (*KLRD1, GZMH, GZMA, NKG7* and *CD52*), CD4^+^ T cells (*BCL6, CD3D, CD4, CD84* and *IL6R*), T-memory cells (*TRBC2, LCK, L7R* and *SELL*), granulocytes (*LYZ*), a minor number of oligodendrocytes and oligodendrocyte-progenitor cell (OPC’s) (*OLIG1, MBP, PDGFA*), and endothelial cells (*CD34, PCAM1, VEGFA*) **Figure 1b, Supplementary Figure 1b-f**.To identify malignant cells, we inferred large scale copy number variations (CNVs) from scRNA-seq profiles by averaging expression over stretches of 100 genes on their respective chromosomes^17^. With this approach, we confirmed that there was minimal contamination by tumor cells (clustered as OPC cells), based on their typical chromosomal alterations (gain in chromosome 7 and loss in chromosome 10), **Supplementary Figure 2a.**

**Figure 1:**
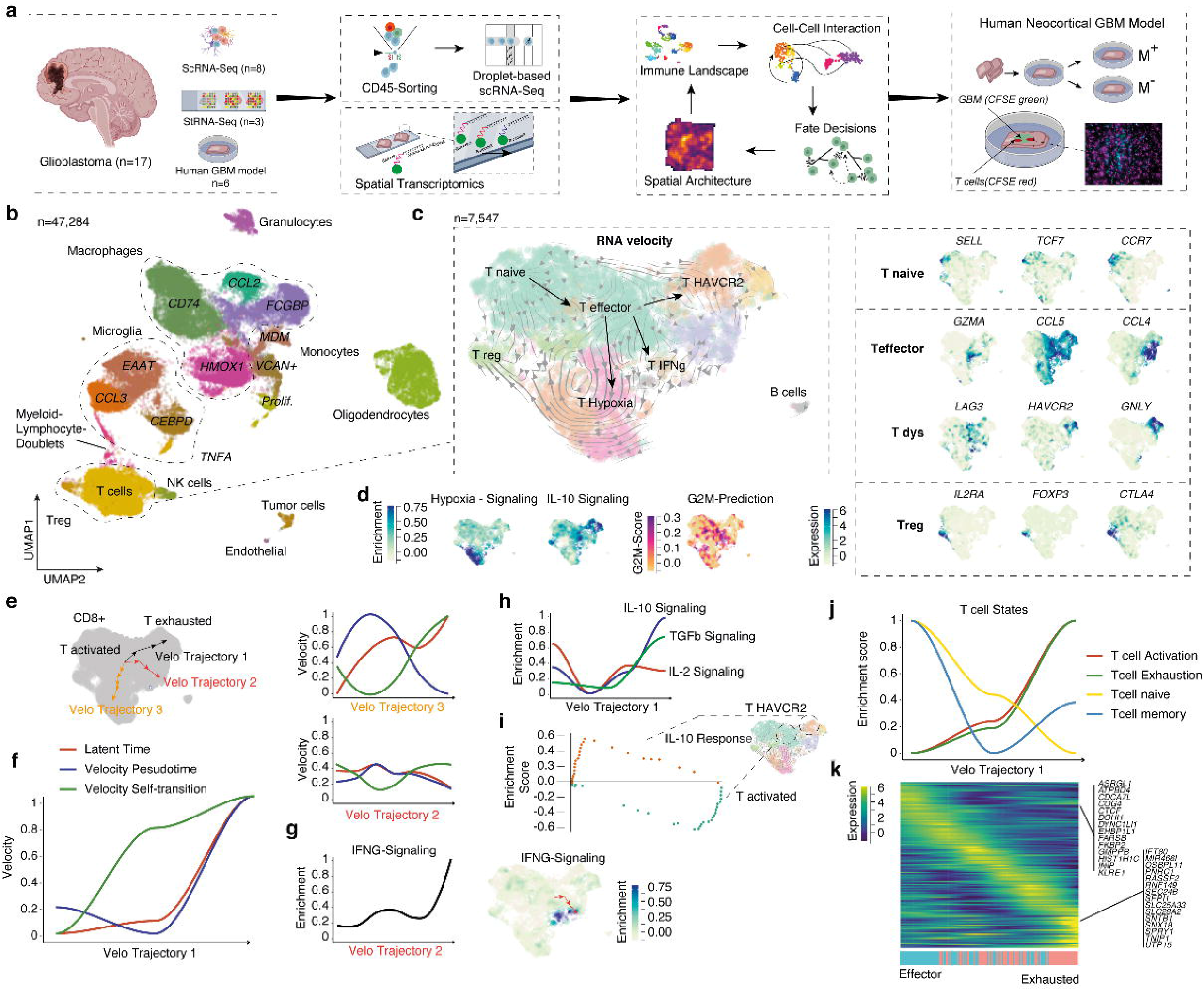
a) Illustration of the workflow, tissue specimens were obtained from 11 glioblastoma patients. Samples from 8 patients were used for scRNA-seq and 3 for spatial transcriptomics. b) Dimensional reduction using UMAP, cell type was determined by SingleR (github.com/dviraran/SingleR). c) Dimensional reduction (UMAP) of CD3+/CD8+ cells. SNN-clustering reveal 13 different clusters. RNA-velocity based pseudotime is presented by streams obtained from dynamic modeling in scvelo. Marker expression in different subpopulations ranked by grade of activity. d) Dimensional reduction plots of estimated gene-set enrichment (left), of cell proliferation (middle) and regulatory marker genes (right). e,f) Detailed presentation of 3 major trajectories and its changes of estimated pseudotime across different models. Velocity trajectory 2 reveals almost no temporal changes, but is marked by increasing IFNg signaling along the trajectory. g) Line plot of IFNg signalling gene expression along the velocity trajectory 2 (left) and dimensional reduction (right). h) Inferring dynamic alterations along the trajectory revealed up-regulation of IL-10 and TGF-ß signaling, presented as a line plot (top). (i) Gene set enrichment analysis of the IL10 signaling enrichment in defined start and destination region of the trajectory (bottom). j) Mapping of gene expression along the pseudotime trajectory. Bars at the bottom indicate the enrichment of each cell to enrich effector or exhausted signatures. Line plots on the top, showed enrichment for the T cell state signatures^8^.

### Diversity of T cells in the Glioblastoma Microenvironment

To investigate the diversity of the T cells present in the microenvironment, we examined them by two different but complementary methods. Firstly, T cells were isolated *in-silico* by means of clustering (as shown above), based on previously published marker gene expression profiles (CD3^+^, CD4^+^/CD8^+^). Secondly, they were isolated using the SCINA model, which resulted in a total of 7,547 cells **Supplementary Figure 1g**. Focusing on the different regulatory states of these cells, we identified 13 subclusters using SNN-clustering which were then re-embedded into a dynamic model using RNA-velocity, closely reflecting different activation states, **Supplementary Figure 1h and Figure 1c**. Reconstruction of lineage differentiation trajectories by means of both pseudotime and latent time provided insights into the transformation of the cells over time^18^, **Figure 1c**. Based on common marker signatures, we defined activated T cells by expressing *GZMA, CCL5* and *CCL4, IL7RI* and other markers (**Supplementary Figure 1i and 2b**) as well as increased proliferation (G2M-score, **Figure 1d**) (C1,C10), and naive T cells by the expression of *SELL*, *TCF7* and *CCR7* (C0), **Figure 1c and Supplementary Figure 1h**. We further identified T cell subgroups marked by hypoxia and heat-shock signaling (*HSPA8*, C2, C5,C6) and by a subgroup which showed mixed expression of activation/dysfunctional/exhaustion markers (*HAVCR2, GNLY*) and strong enrichment IL-10 signaling, **Figure 1d**. We estimated the G2M-score and identified a lower frequency of cell cycle in the differentiated/later states of T cells with the exception of the IFN-gamma subtype, **Figure 1d**. Additional marker plots are given in the **Supplementary Figure 2b**. Regulatory CD4+ T cells (*FOXP3*, *IL2RA* and *CTLA4*) represents a minor population in the glioblastoma microenvironment, **Figure 1c** and **Supplementary Figure 1i-j**.

### Dysfunctional State of T cells is Driven by IL-10 Signaling

To gain insights into the regulatory mechanism of immune cells, we reconstructed fate decisions made during T cell exhaustion using pseudotime trajectories along the estimated velocity streams, **Figure 1e**. Using RNA-velocity, we estimated cells with high probability of initial and terminal states which resulted in 3 major branches with unique cell fate drivers, **Figure 1e**. In order to determine the dynamic adaptation across all branches (Velo trajectory 1-3), we computed pseudotime vectors using various models. Only two trajectories confirmed the predicted ascending pseudotime inference across multiple models (HAVCR2(+)-T cells and hypoxia-induced T cells), **Figure 1f**. We assume that terminal states without significant pseudotime connection do not inevitably arise from the determined initial state, which suggests that the lineage of IFN-gamma driven T cells coexists and most likely originated from an earlier lineage branch **Figure 1g**.

Along our trajectory from effector to HAVCR2(+)T cells, we showed a latent time-dependent increase of the response to anti-inflammatory signaling such as IL-10 and TGF-beta signaling **Figure 1h**. Next, we mapped the gene expression of defined exhausted and effector signatures along our differentiation trajectory, revealing an enrichment of exhausted genes within the destination cluster. Thus, our data suggests that this response to IL-10 contributes to the dysfunctional state of T cells and affects fate decisions **Figure 1i**. To gain further insights into accurate downstream signaling of IL-10, IFN-gamma, and IL2, we created a library of the 50 most highly up-and downregulated genes, **Supplementary Figure 3a-b**. We then extracted signatures observed within the different T cell clusters and compared them with stimulated T cells. As expected, genes upregulated by IL2 stimulation were significantly enriched in cluster 1, while signature genes from IFN-gamma stimulation was enriched in cluster 12. Signature genes from IL-10 stimulation showed a significant enrichment in the dysfunctional cluster (Cluster 9), **Supplementary Figure 3c**. Furthermore, we mapped our signatures along our defined velocity trajectory 1 which confirmed our predicted pathway inference, **Figure 1j-k and Supplementary Figure 3d**.

### T cell Activation and Exhaustion Reveals Spatial Heterogeneity and Association with Glioblastoma Subtypes

Glioblastomas present a high degree of heterogeneity due to regional metabolic differences and varying composition of the tumor microenvironment. Mapping of spatially resolved gene expression is a novel technique which will help overcome the limitations of scRNA-seq, where spatial information is lost. We performed spatial transcriptomic RNA sequencing (stRNA-seq) of 3 primary IDH1/2 wildtype glioblastoma, containing a total number of 2,352 spots, **Figure 2a**. We observed a median of 8 cells per spot (range: 4 to 22 cells per spot), which allows the spatial mapping of gene expression, but not at single cell resolution. However, when we compared our dataset to the latest classification of glioblastoma^19^, consistent results were obtained in accordance with the diversity of subtype expression, **Figure 2b**. In particular, Neftel and colleagues raised evidence that the mesenchymal gene expression is more likely associated with immune response and cellular interaction to myeloid cells^19^. In order to investigate the spatial distribution of the mesenchymal subgroup, we performed gene set enrichment analysis at spatial resolution which revealed spot-wise enrichments within all samples, **Figure 2c-d**. In a next step, we used a seeded non-negative matrix factorization (NMF) regression^20^ to estimate the probability of individual T cell clusters at spatial resolution, **Figure 2e**. We computed the spatial overlap of T cell clusters and glioblastoma states, using a Bayesian approach, which revealed a high estimated correlation of the mesenchymal and T cell cluster 3 (T HAVCR2), cluster 5-7 (T Hypoxia), **Figure 2f**. Further analysis confirming the strong overlap of dysfunctional/exhausted marker (LAG3 and HAVCR2) exclusively with the mesenchymal gene expression signature, **Figure 2g**, suggesting that our data support the findings from Neftel and colleagues.

**Figure 2:**
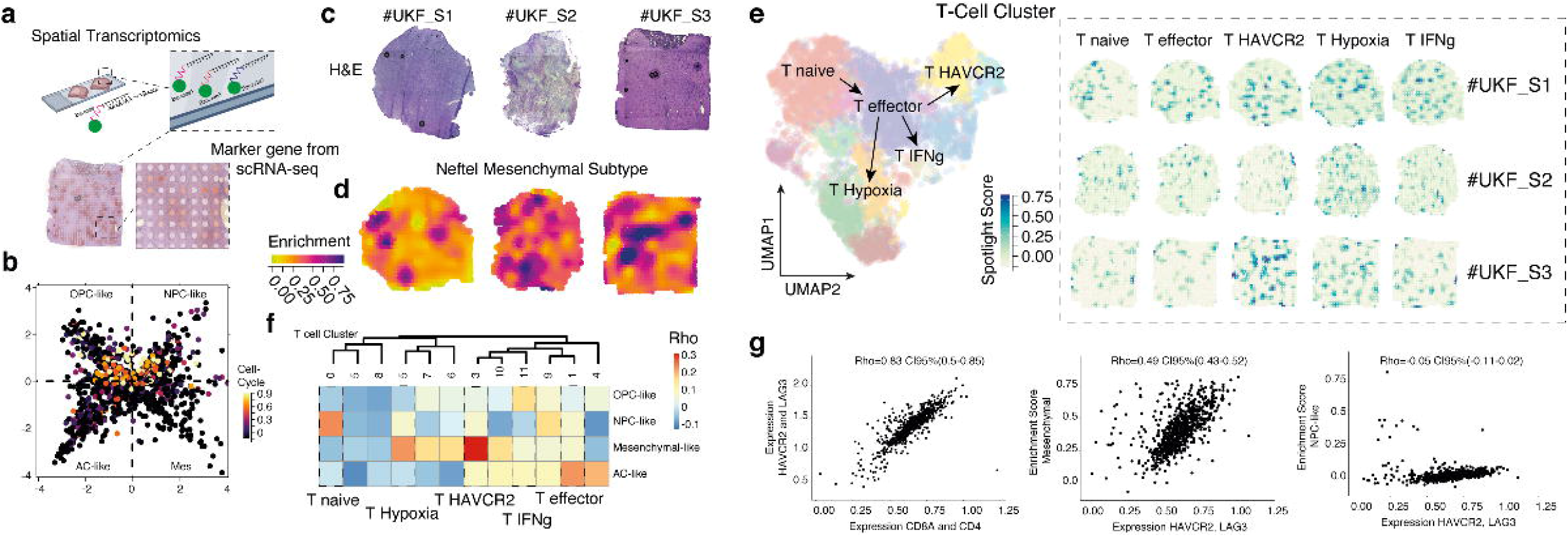
**a)** Workflow of spatial transcriptomics **b)** 2D representation of heterogeneous states in glioblastoma by Neftel, colors indicate the expression of cycling cells (quantile). **c)** H&E stainings and **(d)** gene expression enrichment maps of the mesenchymal state defined by Neftel^19^. **e)** Enrichment maps of T cell clusters by spotlight algorithm^20^ in different samples. Maps are colored by a normalized spotlight score. **f)** Correlation heatmap of a Bayesian correlation model of T cell clusters and glioblastoma transcriptional subtypes. **(g)** Scatter plots of spot-wise correlation of *HAVCR2* and *LAG3* expression and T cells (left side), mesenchymal-like gene expression (middle) and NPC-like gene expression (right side), Rho correlation and CI95% are given at the top.

### A Subset of Microglia and Macrophages Drive IL-10 Stimulation

In our recent investigation^21^, the crosstalk between microglia cells and reactive astrocytes in the tumor microenvironment was found to be responsible for upregulating IL10 release. This is mediated by microglia/macrophages stimulated with IFN gamma, leading to JAK/STAT activation in tumor-associated astrocytes. In this study we introduce the “nearest functionally connected neighbor” algorithm (NFCN), an in-silico model to identify the most likely related cell pairs through divergent down-and up-stream signal activity, **Figure 3a**. In our model, we assume that cellular interaction with distinct mutual activation implies two fundamental prerequisites. On the one hand, the ligand needs to be expressed and released, or otherwise exposed on the cell surface. To avoid the chances of randomly elevated expression or technical artifacts, we also looked at the simultaneous occurrence of ligand induction (upstream pathway signaling). On the other hand, the receptor needs to be expressed and, additionally, downstream signaling has to be activated as well. This allows us to predict the functional status of the receiver cell (Explanation of the model can be found in the Methods section, with an overview in **Supplementary Figure 4)**. We used our *in-silico* model to screen for potential cells responsible for IL-10 activation of T cells. The algorithm identified pairs of lymphoid (T cell clusters) and myeloid cells (macrophages and microglia cluster) and estimated the likelihood of mutual activation **Figure 3b,c**. By extraction of the nearest connected cells (top 1% ranked cells), we identified a subset of myeloid cells characterized by remarkably high *IL10* expression. Most of the receiver cells in the connected cells (top 1% ranked cells) originated from the T cell cluster **Figure 3d**. Our predicted IL10 interactions were also supported by another computational approach (**Supplementary Figure 5).** In order to validate our computational model, we used SPOTlight^20^, an algorithm to predict the spatial position of cells from scRNA-seq data, and were able to confirm a significant overlap. In order to explore the difference between connected and non-connected cells, we extracted both connected and non-connected cells, defined by the highest and lowest interaction-scores (quantile 97.5%) Using differential gene expression analysis, we observed multiple genes which confirmed the non-inflammatory polarization status of highly connected cells.

**Figure 3:**
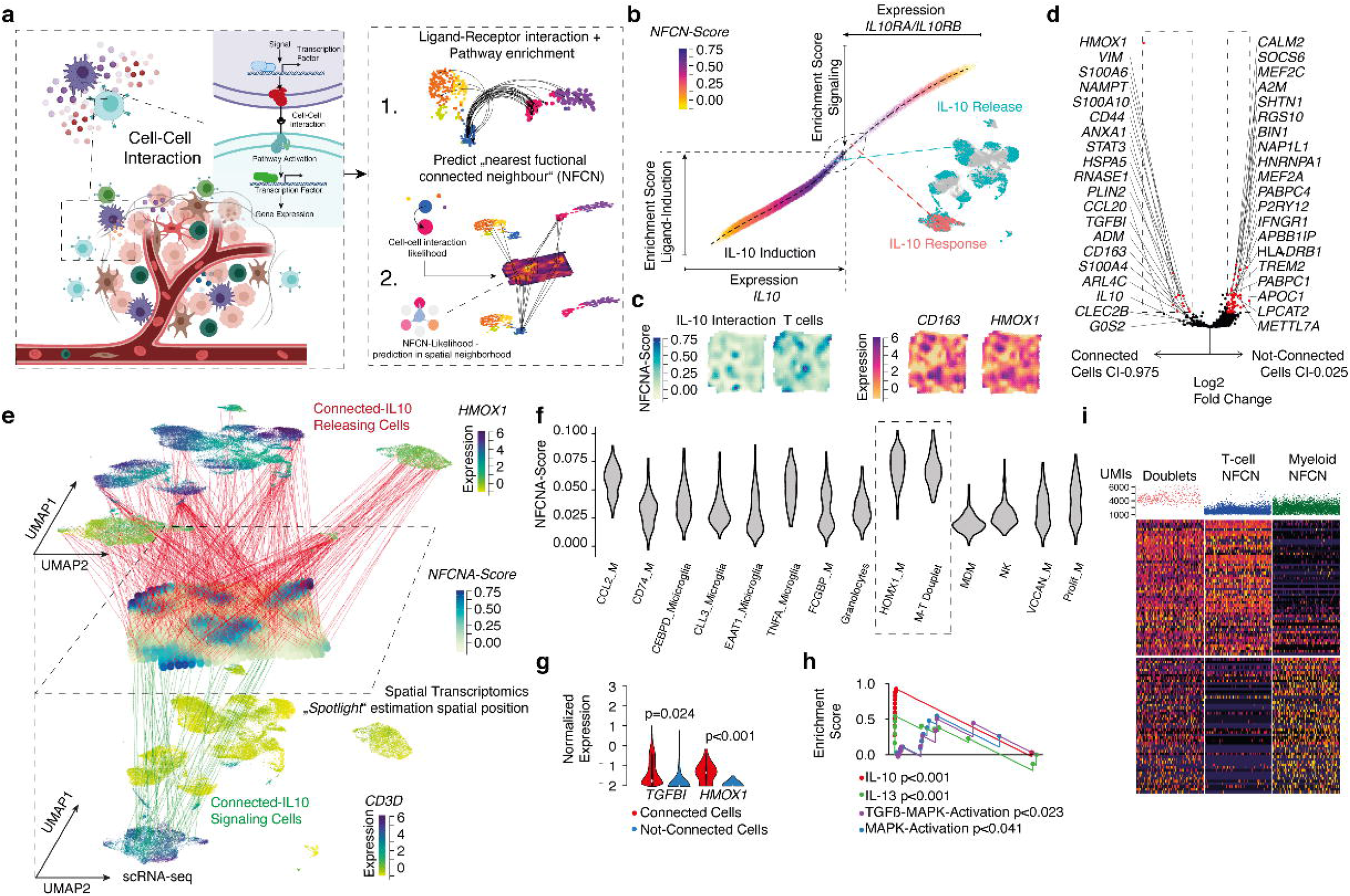
a) Workflow exploring cell-cell interactions using two approaches: 1. Predict functional neighbors based on defined ligand-receptor interaction. 2. Validation in spatial transcriptomic datasets. b) Cell-cell interaction plot as explained in supplementary figure 4. Cells with an interaction-score above 0.95 are mapped to the UMAP. C) Volcano plot of differential gene expression between highly connected cells (CI>97.5%, left side) vs non-connected cells (CI<;2.5%, right side), adjusted −log(p-vale) (FDR) was used at the y-axis. Red cells are defined by fold-change above 2 and FDR < 0.05. d) Spatial surface plots of gene expression and pathway enrichment. e) Multilayer representation of cellular interaction. At the top layer, a UMAP is shown colored by expression levels of HMOX1. Each red line represents a cell (upper CI>99% NFCNA-Score) and its most likely position within the spatial dataset. The second layer is a spatial transcriptomic dataset, colored by the predicted NFCNA-score. At the bottom, the third layer represents a UMAP colored accordingly to CD3 expression. Green lines showed the upper CI>99% of receiver cells. f) Violin plots of the mean NFCNA score in each cluster. g) Violin plots of gene expression between connected cells (CI>97.5%, left side) vs non-connected cells (CI<2.5%, right side). Wilcoxon Rank Sum test and FDR adjustment was used for statistical testing. h) Gene Set enrichment analysis of four different gene sets. i) Heatmap of differently expressed genes or connected myeloid (right) and lymphoid (middle) cells and the cluster of doublets. At the top, scatterplot of total UMIs per cell in each group.

These findings are not surprising, since one of the essential markers of non-inflammatory myeloid cells is *IL10***, Figure 3e**. We showed that the subset of most highly connected cells were marked by *CD163 and* heme oxygenase 1 (*HMOX1)* expression. In a multilayer representation, we illustrated the estimated cellular connections with respect to their most likely spatial coordinates, **Figure 3f**. *HMOX1* is activated during inflammation and oxidative injuries and is regulated through the Nrf2/Bach1-axis, as well as through the IL10/HMOX1-axis. This gene is also well known to be upregulated in the alternative activated macrophage subtype^22^. Another immunosuppressive signaling marker closely related to the alternative activation of macrophages/microglia is the release of TGF-ß^23^, which was also found to be up-regulated in highly connected cells, **Figure 3g**. Consistent with our findings, most downstream signals of the IL10/HMOX1-axis such as STAT3 and p38 MAPK were found to be upregulated in a gene set enrichment analysis, **Figure 3d,h**. In a recent investigation, the role of doublets in the detection of cell-cell interactions was described^24^. In our dataset the score of potential cell-cell connections was similar to HMOX1 marked macrophages, within the doublet dominated cluster. We computed the marker genes of our predicted connected myeloid and lymphoid genes and found each of the marker sets to be highly expressed in the doublet cluster, which further validated the results from our computational model, **Figure 3i**.

### Loss of Myeloid Cells Increases Antitumor Immunity

To provide additional evidence for our findings from the presented computational approach, we made use of the recently described human neocortical GBM model, where the cellular architecture of the CNS is well preserved ^21,25^. We cultured non-infiltrated neocortical slices (defined in a recent report^21,25^) coupled with autografted T cells along with myeloid cell depletion, to understand the communication between myeloid cells in the tumor microenvironment along with lymphoid cells. Three days after chemical depletion of the myeloid cells, we injected a primary cell line (BTSC#233, GFP-tagged, previously characterized by RNA-seq profiling as mesenchymal)^26^. After 4 days of culture, peripheral T cells (same donors), tagged using CellTrace™ Far Red (CTFR) were additionally inoculated and the sections were further cultured for another 48h, **Figure 4a**. Immunostainings showed that myeloid cell depletion reduces the number of IBA1^+^HMOX1^+^ cells, **Figure 4b**. Using an enzyme-linked immunosorbent assay (ELISA) we found a significant reduction in IL10 when the myeloid cells were depleted, regardless of the presence of tumor cells. The strongest difference in IL10 release was observed in myeloid cell depleted sections in the presence of tumor cells, **Figure 4c**. Furthermore, we stained for Granzyme B (GZMB^+^) T cells and quantified IL2 release to examine the amount of effector T cells in both the depleted and non-depleted sections. We found an increased number of GZMB^+^ T cells in sections with myeloid cell depletion, **Figure 4d**, along with a significant increase in IL2 and no differences in IFN gamma release were observed, **Figure 4e-f**. We also stained for the exhaustion marker TIM3 (Gen: *HAVCR2*), which was found to be enriched in T cells in the presence of myeloid cell **Figure 4g**, suggesting that myeloid derived IL10 release (by HMOX1^+^ cells) leads to T cell exhaustion, which is in agreement with the computational model.

**Figure 4:**
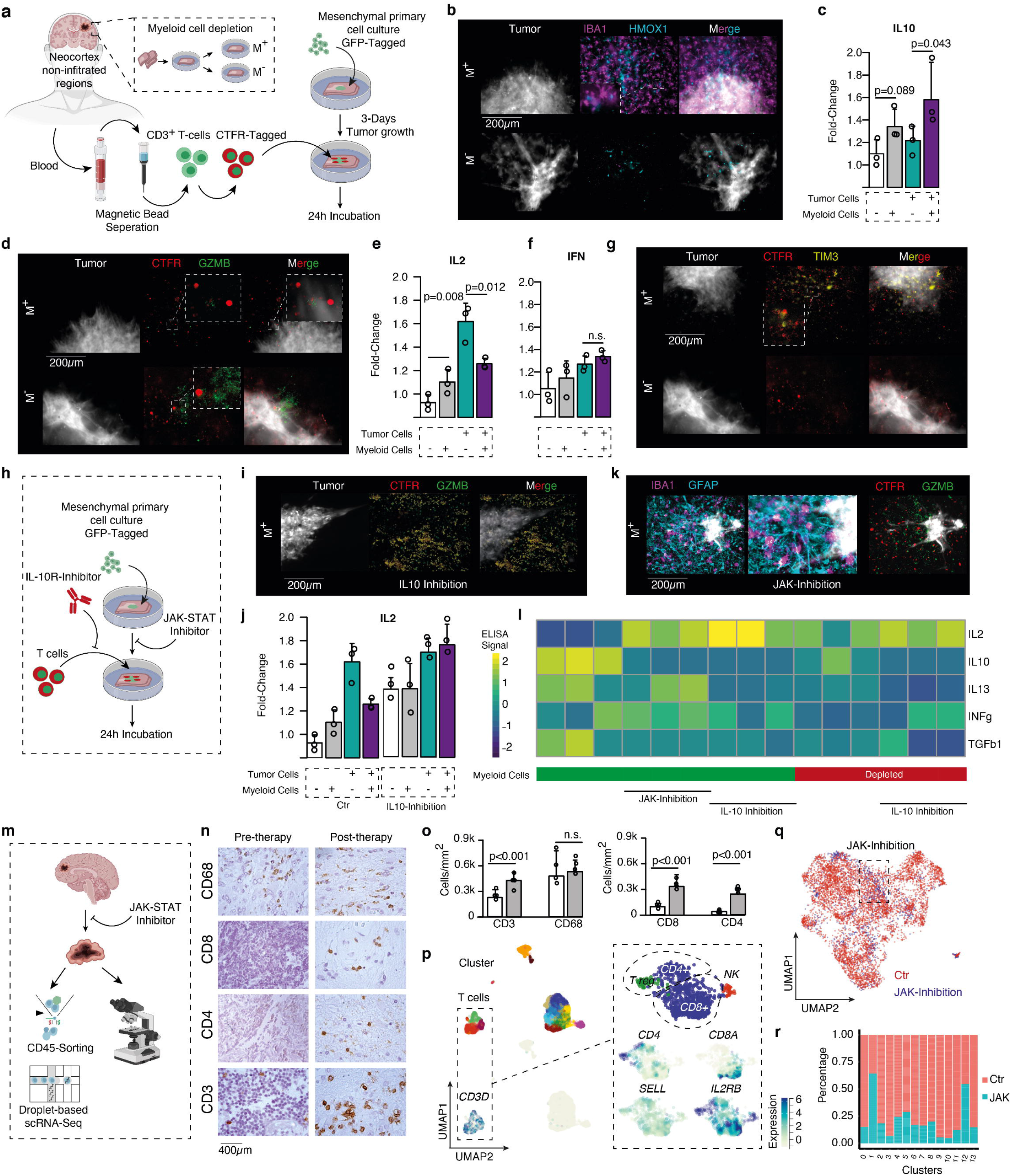
a) Experimental workflow of the neocortical GBM model autografted with patient derived T cells and with/without myeloid cell depletion. b) Immunostainings of IBA1 (Macrophages and Microglia) in magenta and HMOX1 in cyan, tumor cells are depicted in grey. In the upper panel, the control set with no myeloid cell depletion (M^+^) is shown, the bottom panel contains the myeloid cell depleted sections. c) ELISA measurements of IL10 d) Immunostainings of T cells (CSFE-Tagged, in red) and GZMB, a marker of T cell activation (green).). e, f) ELISA measurements of IL2 and IFNg g) Immunostainings of TIM3 (gene: *HAVCR2*) in yellow, which was identified in the scRNA-seq, and T cell in red. h) Illustration of the workflow. i) Immunostainings of tumor cells (grey), T cells (CSFE-Tagged, in red) and GZMB a marker of T cell activation (green) with pre-treatment in anti-IL-10R antibodies. j) ELISA measurements of IL2 in ctr and IL10R inhibition. k) Immunostainings of tumor cells (grey), T cells (CSFE-Tagged, in red) and GZMB a marker of T cell activation (green) treated with JAK-inhibitor Ruxolitinib. l) Heatmap of interleukin intensities different environment. Information regarding the presence of myeloid cells are given at the bottom. m) Illustration of the workflow of JAK inhibition in a recurrent GBM patient. n-o) Immunohistochemistry of immune marker and its quantification, white: pre therapy, gray: post therapy p) Dimensional reduction of single-cell RNA-sequencing of the Ruxolitinib treated patient revealed a large percentage of T cells (right side). q) Comparison of T cells from the Ruxolitinib treated patient and the non-treated cohort. r) A barplot that indicates the cluster specific enrichment of JAK treated T cells. P-values are determined by one-way ANOVA (c,e,f,j) adjusted by Benjamini-Hochberger (c,e,f,j) for multiple testing. Data is given as mean ± standard deviation.

In order to prove that the IL10 signaling is responsible for the induction of the expression of exhausted T cell markers, we preincubated T cells using an IL10 neutralizing inhibitory antibody **Figure 4h**. This resulted in a strong increase of GZMB^+^ T cells, followed by a significant increase of IL2, suggesting that IL10 inhibition drives T cell activity, **Figure 4i-j**.

From our recent study, we found that a JAK/STAT inhibition causing a significant reduction of IL10 within the tumor microenvironment^21^. Based on this investigation, we pre-treated tissue sections with Ruxolitinib, an FDA-approved JAK-inhibitor, before inoculating the sections with patient-derived T cells. We were able to confirm that JAK inhibition caused a significant decrease of IL10 and increased levels of the inflammatory marker IL2, **Figure 4k-l**.

Based on our above findings, we treated a first patient with a recurrent glioblastoma in a neoadjuvant setting with Ruxolitinib for 4 weeks. After resection, we sorted CD45^+^ cells and performed scRNA-sequencing **Figure 4m**. Immunostainings of the tumor revealed a relative increase of CD8^+^ and CD4^+^ T cells whereas CD68^+^ myeloid cells remained stable, **Figure 4n-o**. scRNA sequencing revealed a large number of T cells, most of which express markers for T cell activation and a few cells showing a naive signature, **Figure 4p**. When compared with T cells from our initial dataset, an enrichment of the T cells from the JAK treated patient was seen within cluster 1 (activated T cells) and cluster 12 (B cells), suggesting that JAK-inhibition could be a potential treatment option to boost T cell activation by reducing immunosuppressive programs in both myeloid and glial cells, **Figure 4q-r**.

## Discussion

Although single-cell RNA-sequencing accurately maps the cellular architecture and reflects the diversity of cellular states^17,19,27,28^, there is a lack of spatial information. Here, we combine single cell RNA sequencing of the immune compartment along with spatial transcriptomic RNA-sequencing (stRNA-seq) to gain better insights into the complex crosstalk, cellular states and cellular plasticity leading to the immunosuppressive environment found in glioblastoma (GBM).

Recent studies have reported different subtypes of microglia and macrophages occupying glial tumors^1,4,19,27,28^. However, detailed information about lymphoid infiltration cells is lacking. There is intense interest in T cells and their varied states due to their importance in the development of targeted therapies and to further the understanding of the immunosuppressive environment of glioblastoma. T cell states, particularly in disease, are somehow difficult to accurately classify, leading to numerous definitions and markers in recent years^2,7,29–31^. Some authors use the terms "dysfunctional" and "exhausted" synonymously^32^, whereas others differentiate between the dysfunctional and the exhausted states of T cells^29,31^. In this study we use the definition of cellular states proposed by Singer et al., 2016^8^. On the basis of these gene sets, our data showed that only cells which remained activated along the pseudotime-trajectory were able to enter a state of dysfunction, and later exhaustion. The dysfunction appears to be a transient state, associated with increased proliferation, despite immunosuppressive stimulation from the tumor environment. This imbalance between pro- and anti-inflammatory signaling, dominated by IL10 secretion, leads to final exhaustion of the T cells, which is in agreement with the current literature^2,33^. In order to find a consensus with regard to marker genes we further validated our findings on a set of exhausted marker genes recently published in an overview study^34^. We and others have shown that the GBM microenvironment aids in the evolution of immune suppression. In this process, astrocytes and myeloid cells, both driven by STAT-3 signaling, orchestrate the immunosuppressive environment^4,21,35,36^. Based on the knowledge that IL10 interaction plays a crucial role in the shift from activated to exhausted T cells, we built an *in-silico* model that identified potential connected cells driving T cell exhaustion.

Using this model, we identified a subset of myeloid cells marked by high expression of *HMOX1^+^*, a gene which is induced by oxidative stress and metabolic imbalance^37,38^. *HMOX1* is linked to the STAT-3 pathway and induces IL10 production via MAPK activation, and all of these markers were also found to be upregulated in our connected cells, as reported above. Furthermore, we used spatial transcriptomics to confirm the spatial overlap of cells that we identified as highly connected. We were able to show that the *HMOX1^+^*-myeloid cells were spatially correlated with exhaustion and the mesenchymal state of glioblastoma. These findings are in accord with published reports, revealing that the mesenchymal cells are the component of GBM responsible for the immune crosstalk^19^. HMOX1 expression in GBM and IDH-WT astrocytoma was found to be increased in recurrent GBM and negatively associated with overall survival, **Supplementary Figure 6a,b**. In addition, we made use of a human neocortical GBM model coupled with patient derived T cells in addition to depletion of myeloid cells. This model helped us to simulate the function of the myeloid cells with regard to IL10 release and T cell stimulation. Fitting with our computational model, we confirmed that HMOX1^+^ myeloid cells cause a reduction of effector T cells, with a respective reduction in IL2 release and increased expression of our identified exhaustion marker TIM3. Following our recent investigations in which we demonstrated that JAK-inhibition is able to reduce the level of IL10 in human brain tumors^21^, we demonstrated that in a single patient that inhibition of the JAK-STAT axis was able to partially rescue the immunosuppressive environment. The treated subject is still alive but showed permanent disruption of the blood-brain barrier with repetitive increase of contrast enhancing lesions. Each radiologically confirmed progress was sampled without evidence of a tumor recurrence, suggesting that manipulation of the glia/myeloid environment simultaneously caused exaggerated inflammation and pseudo progression. Our single-cell RNA-seq confirmed a pronounced enrichment of activated T cells, while the number of myeloid cells remained relatively stable. In this sample, we also detected the strongest contamination with glial cells. We assumed that potential doublets or strong cell-cell connections led to glial cells being detected as CD45^+^, resulting in a false positive sorting.

In conclusion, this work provides the first knowledge regarding lymphocyte population in the glioblastoma microenvironment where we showed that the functional interaction between myeloid and lymphoid cells, leads to a dysfunctional state of T cells. Using human neocortical GBM model and single patient subject we showed that the IL-10 driven T cell exhaustion can be rescued by JAK/STAT inhibition. Thus, the results from this work can be the steppingstone towards successful immunotherapeutic approaches for GBM.

## Methods

### Ethical Approval

The local ethics committee of the University of Freiburg approved the data evaluation, imaging procedures and experimental design (protocol 100020/09 and 472/15_160880). The methods were carried out in accordance with the approved guidelines, with written informed consent obtained from all subjects. The studies were approved by an institutional review board. Further information and requests for resources, raw data and reagents should be directed and will be fulfilled by the Contact: D. H. Heiland. A complete table of all materials used is given in the supplementary information.

### T cell isolation and stimulation

Blood was drawn from a healthy human individual into an EDTA (ethylenediaminetetraacetic acid) cannula. T cells were extracted in a negative selection manner using a MACSxpress® Whole Blood Pan T Cell Isolation Kit (Miltenyi Biotech). T cells were then transferred in Advanced RPMI 1640 Medium (ThermoFisher Scientific, Pinneberg, Germany) and split for cytokine treatment: Three technical replicates were used for each T cell-treatment condition. Interleukin 2 (IL-2, Abcam, Cambridge, UK) was used at a final concentration of 1 ng/ml, Interleukin 10 (IL-10, Abcam) at 5 ng/ml, Interferon gamma (IFN-γ, Abcam) at 1 ng/ml and Osteopontin (SPP-1, Abcam) at 3 μg/ml. Cytokine treatment was performed in Advanced RPMI 1640 Medium and T cells were incubated at 37°C and 5% CO2 for 24h.

### RNA sequencing of stimulated T Cells

The purification of mRNA from total RNA samples was achieved using the Dynabeads mRNA Purification Kit (Thermo Fisher Scientific, Carlsbad, USA). The subsequent reverse transcription reaction was performed using SuperScript IV reverse transcriptase (Thermo Fisher Scientific, Carlsbad, USA). For preparation of RNA sequencing, the Low Input by PCR Barcoding Kit and the cDNA-PCR Sequencing Kit (Oxford Nanopore Technologies, Oxford, United Kingdom) were used as recommended by the manufacturer. RNA sequencing was performed using the MinION Sequencing Device, the SpotON Flow Cell and MinKNOW software (Oxford Nanopore Technologies, Oxford, United Kingdom) according to the manufacturer’s instructions. Samples were sequenced for 48h on two flow-cells. Basecalling was performed by Albacore implemented in the nanopore software. Only D^2^-Reads with a quality Score above 8 were used for further alignment.

### Sequence trimming and Alignment

In the framework of this study, we developed an automated pipeline for nanopore cDNA-seq data, which is available at github (https://github.com/heilandd/NanoPoreSeq). First the pipeline set up a new class termed “Poreseq” by a distinct sample description file. The analysis starts by rearranging the reads from the *fastq* output from the nanopore sequencer containing all of the D^2^-Reads. All *fastq* files need to be combined into one file. Multiplexed samples were separated according to their barcode and trimmed by Porechop (https://github.com/rrwick/Porechop). Alignment was performed with minimap2 (https://github.com/lh3-/minimap2) and processed with sam-tools.

### Posthoc Analysis of Bulk-RNA-seq

A matrix of counted genes was further prepared by the RawToVis.R (github.com/heilandd/VRSD_Lab_v1.5) script, containing normalization of Mapped reads by DESeq, batch effect removal (ComBat package) and fitting for differential gene expression. Gene set enrichment analysis was performed by transformation of the log2 fold-change of DE into a ranked z-scored matrix, which was used as the input. The expression matrix was analysed with AutoPipe (https://github.com/heilandd/AutoPipe) by a supervised machine-learning algorithm and visualized with a heatmap. Full analysis was visualized with the Visualization of RNA-Seq Data (VRSD_Lab software, github.com/heilandd/VRSD_Lab_v1.5) as a dashboard app based on shiny R-software. We extracted the 50 top up/down regulated genes respectively of each stimulation with respect to control condition to construct a stimulation library.

### Single-Cell Suspension for scRNA-sequencing

Tumor tissue was obtained from glioma surgery immediately after resection and was transported in phosphate-buffered saline (PBS) within approximately 5 minutes into our cell culture laboratory. Tumor tissue was processed under a laminar flow cabinet. Tissue was reduced to small pieces using two scalpels and the tissue was processed with the Neural Tissue Dissociation Kit (T) using C-Tubes (Miltenyi Biotech, Bergisch-Gladbach, Germany) according to the manufacturer’s instructions. The Debris Removal Kit from Miltenyi was used according to the manufacturer’s instructions to remove remaining myelin and extracellular debris. In order to remove the remaining erythrocytes, we resuspended the pellet in 3,5 ml ACK lysis buffer (ThermoFisher Scientific, Pinneberg, Germany) and incubated the suspension for 5 minutes followed by a centrifugation step (350g, 10 min, RT). Cell quantification with a hematocytometer was performed after discarding the supernatant and resuspending the pellet in PBS. Cell suspensions were centrifuged again (350g, 10 min, RT) and resuspended in freezing medium containing 10% DMSO (Sigma-Aldrich, Schnelldorf, Germany) in FCS (PAN-Biotech, Aidenbach, Germany). Cell suspensions were immediately placed in a freezing box containing isopropanol and stored in a −80°C freezer for not more than 4 weeks.

### Cell sorting by Magnetic Beads

Four frozen single-cell suspensions, originating from one patient with an IDH-mutated glioma and three patients with an IDH-wildtype glioblastoma (GBM), were thawed and the dead cells magnetically labeled and eliminated using a Dead Cell Removal Kit (Miltenyi Biotech). The tumor immune environment in general and T cells in particular were positively selected by using CD3+-MACS (Miltenyi Biotech). Cells were stained with trypan blue, counted using a hematocytometer and prepared at a concentration of 700 cells/μL.

### Droplet scRNA-sequencing

At least 16000 cells per sample were loaded on the Chromium Controller (10x Genomics, Pleasanton, CA, USA) for one reaction of the Chromium Next GEM Single Cell 3’v3.1 protocol (10x Genomics), based on a droplet scRNA-sequencing approach. Library construction and sample indexing was performed according to the manufacturer’s instructions. scRNA-libraries were sequenced on a NextSeq 500/550 High Output Flow Cell v2.5 (150 Cycles) on an Illumina NextSeq 550 (Illumina, San Diego, CA, USA). The bcl2fastq function and the cell ranger (v3.0) was used for quality control.

### Postprocessing scRNA-sequencing

We used cell ranger to detect low-quality read pairs of single-cell RNA sequencing (scRNA-seq) data. We filtered out reads which did not reach the following criteria: (1) bases with quality < 10, (2) no homopolymers (3) ‘N’ bases accounting for ≥10% of the read length. Filtered reads were mapped by STAR aligner and the resulting filtered count matrix further processed by Seurat v3.0 (R-package). We normalized gene expression values by dividing each estimated cell by the total number of transcripts and multiplied by 10,000, followed by natural-log transformation. Next, we removed batch effects and scaled data by a regression model including sample batch and percentage of ribosomal and mitochondrial gene expression. For further analysis we used the 2000 most variable expressed genes and decomposed eigenvalue frequencies of the first 100 principal components and determined the number of non-trivial components by comparison to randomized expression values. The obtained non-trivial components were used for SNN clustering followed by dimensional reduction using the UMAP algorithm. Differently expressed genes (DE) of each cluster were obtained using a hurdle model tailored to scRNA-seq data which is part of the MAST package. Cell types were identified by 3 different methods; Classical expression of signature markers of immune cells; SingleR an automated annotation tool for single-cell RNA sequencing data obtaining signatures from the Human Primary Cell Atlas, SCINA, a semi-supervised cell type identification tool using cell-type signatures as well as a Gene-Set Variation Analysis (GSVA). Results were combined and clusters were assigned to the cell type with the highest enrichment within all models. In order to individually analyze T cells, we used the assigned cluster and filter for the following criteria. For further analysis T cells were defined by: CD3^+^CD8^+^ / CD4^+^CD14^−^LYZ^−^ GFAP^−^CD163^−^IBA^−^.

### Spatial Transcriptomics

The spatial transcriptomics experiments were done using the 10X Spatial transcriptomics kit (https://spatialtranscriptomics.com/). All the instructions for Tissue Optimization and Library preparation were followed according to the manufacturer’s protocol. Here, we briefly describe the methods followed using the library preparation protocol.

### Tissue collection and RNA quality control

Tissue samples from three patients, diagnosed with WHO IV glioblastoma multiforme (GBM), were included in this study. Fresh tissue collected immediately post resection was quickly embedded in optimal cutting temperature compound (OCT, Sakura) and snap frozen in liquid N2. The embedded tissue was stored at −80°C until further processing. A total of 10 sections (10μm each) per sample were lysed using TriZOl (Invitrogen, 15596026) and used to determine RNA integrity. Total RNA was extracted using PicoPure RNA Isolation Kit (Thermo Fisher, KIT0204) according to the manufacturer’s protocol. RIN values were determined using a 2100 Bioanalyzer (RNA 6000 Pico Kit, Agilent) according to the manufacturer’s protocol. It is recommended to only use samples with an RNA integrity value >7.

### Tissue staining and Imaging

Sections were mounted onto spatially barcoded glass slides with poly-T reverse transcription primers, with one section per array. These slides can be stored at −80°C until use. The slides were then warmed to 37°C, after which the sections were fixed for 10 minutes using 4% formaldehyde solution (Carl Roth, P087.1), which was then washed off using PBS. The fixed sections were covered with propan-2-ol (VWR, 20842312). Following evaporation for 40 seconds, sections were incubated in Mayer’s Hematoxylin (VWR, 1092490500) for 7 min, bluing buffer (Dako, CS70230-2) for 90 seconds and finally in Eosin Y (Sigma, E4382) for 1 min. The glass slides were then washed using RNase/DNase free water and incubated at 37°C for 5 min or until dry. Before imaging, the glass slides were mounted with 87% glycerol (AppliChem, A3739) and covered with coverslips (R. Langenbrinck, 01-2450/1). Brightfield imaging was performed at 10x magnification with a Zeiss Axio Imager 2 Microscope, and post-processing was performed using ImageJ software.

The coverslips and glycerol were removed by washing the glass slides in RNase/DNase free water until the coverslips came off, after which the slides were washed using 80% ethanol to remove any remaining glycerol.

### Permeabilization, cDNA synthesis and tissue removal

For each capture array, 70μL of pre-permeabilization buffer, containing 50U/μL Collagenase along with 0.1% Pepsin in HCl was added, followed by an incubation for 20 minutes at 37°C. Each array well was then carefully washed using 100μL 0.1x SSC buffer. 70μL of Pepsin was then added and incubated for 11 minutes at 37°C. Each well was washed as previously described and 75μL of cDNA synthesis master mix containing: 96μL of 5X First strand buffer, 24 μL 0.1M DTT, 255.2μL of DNase/RNase free water, 4.8μL Actinomycin, 4.5μL of 20mg/mL BSA, 24μL of 10mM dNTP, 48μL of Superscript^®^ and 24μL of RNAseOUT™ was added to each well and incubated for 20 hours at 42°C without shaking. Cyanine 3-dCTP was used to aid in the determination of the footprint of the tissue section used.

Since glioblastoma tissue is a fatty tissue, degradation and tissue removal was carried out using Proteinase K treatment for which 420μL Proteinase K and PKD buffer (1:7), were added to each well and then incubated at 56°C for 1hr with intermittent agitation (15 seconds / 3 minutes). After incubation, the glass slides were washed three times with 100mL of 50°C SSC/SDS buffer with agitation for 10 minutes, 1 minute and finally for 1 minute at 300 rpm. The glass slides were then air-dried at room temperature. Tissue cleavage was carried out by the addition of 70μL of cleavage buffer (320μL RNase/DNase free water, 104μL Second strand buffer, 4.2μL of 10mM dNTP, 4.8μL of 20 mg/mL BSA and 48μL of USER™ Enzyme) to each well and incubation at 37°C for 2 hours with intermittent agitation.

### Spot Hybridization

In order to determine the exact location and quality of each of the 1007 spots, fluorescent Cyanine-3 A is hybridized to the 5’ ends of the surface probes. 75μL of the hybridization solution (20μL of 10μM Cynaine-3A probe and 20μL of 10μM Cyanine-3 Frame probe in 960μL of 1X PBS) was added to each well and incubated for 10min at room temperature. The slides were then washed three times with 100ml of SSC/SDS buffer preheated to 50°C for 10min, 1min and 1min at room temperature with agitation. The slides were then air-dried and imaged after applying Slowfade^®^ Gold Antifade medium and a coverslip.

### Library Preparation

#### 1. Second Strand Synthesis

5μL second strand synthesis mix containing 20μL of 5X First Strand Buffer, 14μL of DNA polymerase I (10U/μL) and 3.5μL Ribonuclease H (2U/μL) were added to the cleaved sample and incubated at 16°C for 2 hours. Eppendorf tubes were placed on ice and 5μL of T4 DNA polymerase (3U/μL) were added to each strand and incubated for 20 minutes at 16°C. 25μL of 80mM EDTA (mix 30μL of 500mM EDTA with 158μL DNase/RNase free water) was added to each sample and the samples were kept cool on ice.

#### 2. cDNA purification

cDNA from the previous step was purified using Agencourt RNAclean XP beads and DynaMag™-2 magnetic rack, incubated at room temperature for 5 min. Further cleansing was performed by the addition of 80% Ethanol to the sample tubes, while the samples were still placed in the magnetic rack. Sample elution was then carried out using 13μL of NTP/water mix.

#### 3. *In Vitro* Transcription and Purification

cDNA transcription to aRNA was carried out by adding 4μL of reaction mix containing: 10x Reaction Buffer, T7 Enzyme mix and SUPERaseIn^TM^ RNase Inhibitor (20 U/μL) to 12μL of the eluted cDNA sample and incubated at 37°C, for 14 hours. The samples were purified using RNA clean XP beads according to the manufacturer’s protocol and further eluted into 10μL DNase/RNase free water. The amount and average fragment length of amplified RNA was determined using the RNA 6000 Pico Kit (Agilent, 5067-1513) with a 2100 Bioanalyzer according to the manufacturer’s protocol.

#### 4. Adapter Ligation

Next, 2.5μL Ligation adapter (IDT) was added to the sample and was heated for 2 min at 70°C and then placed on ice. A total of 4.5μL ligation mix containing 11.3μL of 10X T4 RNA Ligase, T4 RNA truncated Ligase 2 and 11.3μL of murine RNase inhibitor was then added to the sample. Samples were then incubated at 25°C for 1 hour. The samples were then purified using RNAClean XP beads according to the manufacturer’s protocol.

#### 5. Second cDNA synthesis

Purified samples were mixed with 1μL cDNA primer (IDT), 1μL dNTP mix up to a total volume of 12μL and incubated at 65°C for 5 min and then directly placed on ice. A 1.5ml Eppendorf tube 8μL of the sample was mixed with 30μL of First Strand Buffer(5X),), 7.5μL of DTT(0.1M), 7.5μL of DNase/RNase free water, 7.5μL of SuperScript® III Reverse transcriptase and 7.5μL of RNaseOUT™ Recombinant ribonuclease Inhibitor and incubated at 50°C for 1 hour followed by cDNA purification using Agencourt RNAClean XP beads according to the manufacturer’s protocol. Samples were then stored at −20°C.

#### 6. PCR amplification

Prior to PCR amplification, we determined that 20 cycles were required for appropriate amplification. A total reaction volume of 25μL containing 2x KAPA mix, 0.04μM PCRInPE2 (IDT), 0.4μM PCR InPE1.0 (IDT), 0.5μM PCR Index (IDT) and 5μL of purified cDNA were amplified using the following protocol: 98°C for 3 min followed by 20 cycles at 98°C for 20 seconds, 60°C for 30 seconds, 72°C for 30 seconds followed by 72°C for 5 minutes. The libraries were purified according to the manufacturer’s protocol and eluted in 20μL EB (elution buffer). The samples were then stored at −20°C until used.

#### 7. Quality control of Libraries

The average length of the prepared libraries was quantified using an Agilent DNA 1000 high sensitivity kit with a 2100 Bioanalyzer. The concentration of the libraries was determined using a Qubit dsDNA HS kit. The libraries were diluted to 4nM, pooled and denatured before sequencing on the Illumina NextSeq platform using paired-end sequencing. We used 30 cycles for read 1 and 270 cycles for read 2 during sequencing.

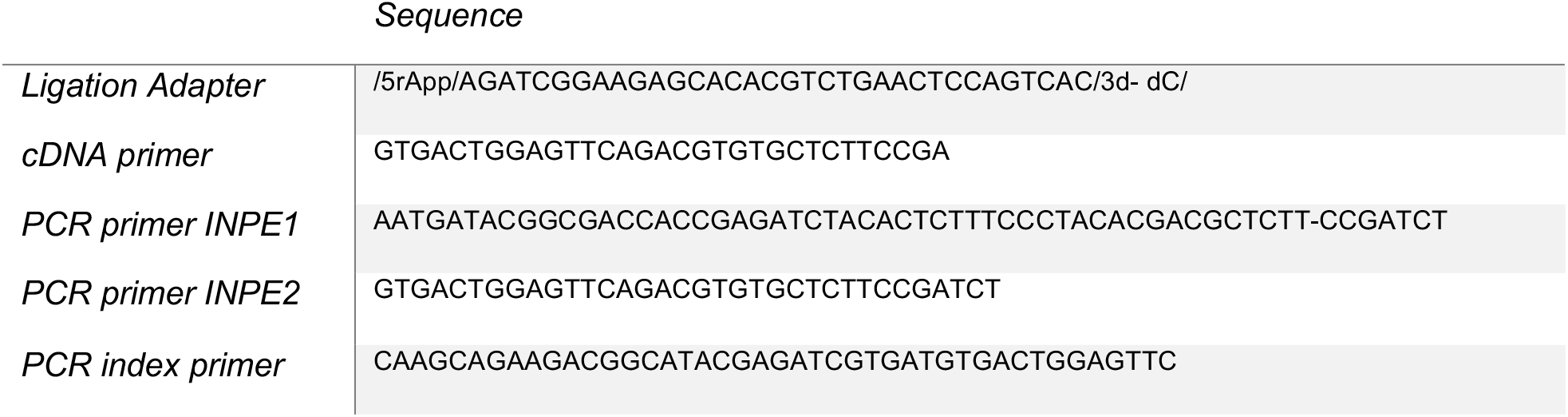

### Postprocessing Spatial Transcriptomics

First, we aligned the H&E staining by the use of the st-pipeline (github.com/SpatialTranscriptomics-Research/st_pipeline). The pipeline contains the following steps: Quality trimming and removing of low quality bases (bases with quality < 10), sanity check (reads same length, reads order, etc‥), remove homopolymers, normalize for AT and GC content, mapping the read2 with STAR, demultiplexing based on read1, sort for reads (read1) with valid barcodes, annotate the reads with htseq-count, group annotated reads by barcode (spot position), gene and genomic location (with an offset) to get a read count (github.com/SpatialTranscriptomics-Research/st_pipeline). The pipeline resulted in a gene count matrix and a spatial information file containing the x and y position and the H&E image. We used the Seurat v3.0 package to normalize gene expression values by dividing each estimated cell by the total number of transcripts and multiplied by 10,000, followed by natural-log transformation. As described for sc-RNA sequencing, we removed batch effects and scaled data by a regression model including sample batch and percentage of ribosomal and mitochondrial gene expression. For further analysis we used the 2000 most variable expressed genes and decomposed eigenvalue frequencies of the first 100 principal components and determined the number of non-trivial components by comparison to randomized expression values. The obtained non-trivial components were used for SNN clustering followed by dimensional reduction using the UMAP algorithm. Differently expressed genes (DE) of each cluster were obtained using a hurdle model tailored to scRNA-seq data which is part of the MAST package. We further build a user-friendly viewer for spatial transcriptomic data and provide tutorials on analysis of data: https://themilolab.github.io/SPATA/index.html.

### Spatial gene expression

For spatial expression plots, we used either normalized and scaled gene expression values (to plot single genes) or scores of a set of genes, using the 0.5 quantile of a probability distribution fitting. The x-axis and y-axis coordinates are given by the input file based on the localization at the H&E staining. We computed a matrix based on the maximum and minimum extension of the spots used (32×33) containing the gene expression or computed scores. Spots without tissue covering were set to zero. Next, we transformed the matrix, using the squared distance between two points divided by a given threshold, implemented in the fields package (R-software) and adapted the input values by increasing the contrast between uncovered spots. The data are illustrated either as surface plots (plotly package R-software) or as images (graphics package R-software).

### Representation of Cellular States

We aligned cells/spots to variable states with regard to gene sets (GS) that were selected GS_(1,2,‥n)_. First, we separated cells into GS_(1+2)_ versus GS_(2+4)_, using the following equation:

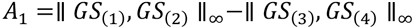

A1 defines the y-axis of the two-dimensional representation. In a next step, we calculated the x-axis separately for spots A1<0 and A1>0:

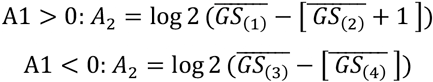

For further visualization of the enrichment of subsets of cells according to gene set enrichment across the two-dimensional representation, we transformed the distribution to representative colors using a probability distribution fitting. This representation is an adapted method published by Neftel and colleges recently^19,28^.

### Spatial correlation analysis

The spatial correlation was performed by integrating a deep autoencoder for background noise reduction and a Bayesian correlation model. In a first step, we performed noise reduction through an autoencoder similar to recent described for single-cell RNA-sequencing studies^39^. The autoencoder consist of an encoder and decoder part which can be defined as transitions:

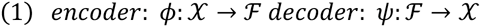

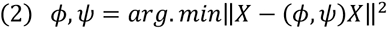

The encoder stage of an autoencoder takes the input 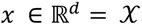 and maps it to 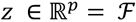 at the layer position φ:

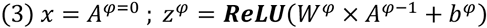

*z*^φ^ is also referred to as *latent representation*, here presented as *z*^1^, *z*^2^,…,*z*^φ = *n*^ in which φ describes the number of hidden layers. **W** is the weight matrix and **b** represent the dropout or bias vector. Our network architecture contained 32 hidden layers, as recommended^39^. In the decoder weights and biases are reconstructed through backpropagation 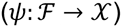 and z is mapped to *x*′ = *A*^0^′ in the shape as *x*′.

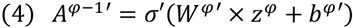

In this context, *W*′, σ′, *b*′ from the decoder are unrelated to *W*, σ, *b* from the encoder. We used a loss function to train the network in order to minimize reconstruction errors.

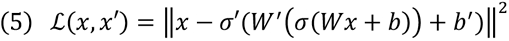

In the second step, we used the predicted gene expression matrix (*x*′) and fitted and Bayesian correlation model (Bayesian First Aid, https://github.com/rasmusab/bayesian_first_aid). An illustration of the spatial correlation is given in the **supplementary figure 7**

### sc-RNA-sequencing integration into spatial context

In order to integrate determined cluster into the spatial context of the spatial transcriptomic data. We used the recent published spotlighted algorithm and integrated the output into our SPATA objects for visualization^20^.

### RNA-velocity and Pseudotime trajectory analysis

In order to determine dynamic gene expression changes, we extracted spliced and unspliced genes from the bam output created by cell ranger using the velocyto.py tool^40^. The resulting *.loom files were merged and transformed into .h5ad format for further processing by scVelo^41^ and CellRank^42^. The pipeline is integrated into the SPATA toolbox^43^. Single-cell data are reformatted into an SPATA S4 object using the UMAP coordinates as spatial coordinates. Outputs of the scVelo script (implemented in the development branch of the SPATA toolbox) are imported into the SPATA S4 object *(slot:@fdata*) and available for visualization. RNA-velocity streams are converted into trajectories and imported to the SPATA S4 object *(slot:@trajectories*). Dynamic gene expression changes along trajectories are performed by the assessTrajectoryTrends() function.

### Gene set enrichment analysis

Gene sets were obtained from the database MSigDB v7 and internally created gene sets are available at githunb.com/heilandd. For enrichment analysis of single clusters, the normalized and centered expression data were used and further transformed to z-scores ranging from 1 to 0. Genes were ranked in accordance to the obtained differential expression values and used as the input for GSEA.

### Identification of cycling cells

We used the set of genes published by Neftel and colleagues to calculate proliferation scores based on the GSVA package implemented in R-software. The analysis based on a non-parametric unsupervised approach, which transformed a classic gene matrix (gene-by-sample) into a gene set by sample matrix resulted in an enrichment score for each sample and pathway. From the output enrichment scores we set a threshold based on distribution fitting to define cycling cells.

### Nearest Functionally Connected Neighbor (NFCN)

To identify connected cells that interact by defined activation or inhibition of down-stream signals in the responder cell, we created a novel model. Therefore, we assumed that a cell-cell interaction is given only if a receptor/ligand pair induce correspondent down-stream signaling within the responder cell (cell with expressed receptor). Furthermore, we take into account that the importance of an activator cell (cell with expressed ligand) can be ranked according to their enriched signaling, which is responsible for inducing ligand expression. Based on these assumptions we defined an algorithm to map cells along an interaction-trajectory. The algorithm was designed to identify potential activators from a defined subset of cells.

As input for the analysis we used a normalized and scaled gene expression matrix, a string containing the subset of target cells, a list of genes defining ligand induction on the one side and receptor signaling on the other side. These genes were chosen either by the MSigDB v7 database or our stimulation library explained above. Next, we down-scaled the data to 3000 representative cells including all myeloid cell types and calculated the enrichment of induction and activation of the receptor/ligand pair. Enrichment scores were calculated by singular value decomposition (SVD) over the genes in the gene set and the coefficients of the first right-singular vector defined the enrichment of induction/activation profile. Both expression values and enrichment scores were fitted by a probability distribution model and cells outside the 95% quantile were removed. Next, we fitted a model using a non-parametric kernel estimation (Gaussian or Cauchy-Kernel), on the basis of receptor/ligand expression (A_exp_) and up/downstream signaling (A_eff_) of each cell (i={1,‥n}). Both input vectors were normalized and z-scored:

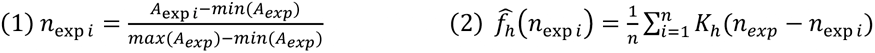

K is the kernel and 0.7 > h > 0.3 is used to adjust the estimator. The model resulted in a trajectory which was defined as Ligand(-)-Induction(-) to cells of the target subset with Receptor(-)-Activation(-).Further cells were aligned along the “interaction-trajectory”. We defined connected cells by reaching the upper 70% CI in receptor/ligand expression as well as sores of induction/activation. The process of representation is illustrated schematically in **Supplementary Figure 4**. Additionally we determined receptor-ligand interaction by the NicheNet software as recommended by the authors^44^.

### CNV estimation

Copy-number Variations (CNVs) were estimated by aligning genes to their chromosomal location and applying a moving average to the relative expression values, with a sliding window of 100 genes within each chromosome, as described recently^17^. First, we arranged genes in accordance to their respective genomic localization using the CONICSmat package (R-software). As a reference set of non-malignant cells, we in-silico extracted 400 CD8 positive cells (unlikely to be expressed on tumor cells). To avoid the considerable impact of any particular gene on the moving average we limited the relative expression values [-2.6,2.6] by replacing all values above/below *exp(i)*=|2.6|, by using the infercnv package (R-software). This was performed only in the context of CNV estimation as previous reported^11^.

### Flow cytometry

Single-Cell suspensions were obtained after Dead-Cell Removal and CD3 MACS-enrichment. Cells were incubated with VivaFix™ 398/550 (BioRad Laboratories, CA, USA) according to the manufacturer’s instructions. Cells were fixed in 4% paraformaldehyde (PFA) for 10 minutes. After centrifugation (350 g; 4°C; 5 min) and removal of the supernatant, the cell pellet was suspended in 0.5 ml 4°C cold FACS buffer. Cell suspensions were washed and centrifuged at 350xg for 5 mins, followed by resuspension in FACS buffer. The washing step was repeated twice. Finally, cells were resuspended in at least 0.5 to 1 mL of FACS buffer depending on the number of cells. We used a Sony SP6800 spectral analyzer in standardization mode with PMT voltage set to maximum to reach a saturation rate below 0.1 %. Gating was performed by FCS Express 7 plus at the Lighthouse Core Facility, University of Freiburg.

### Immunofluorescence

The same protocol was followed for human neocortical slices with or without microglia and tumor cell injection. The media was removed and exchanged for 1 mL of 4% paraformaldehyde (PFA) for 1 h and further incubated in 20% methanol in PBS for 5 minutes. Slices were then permeabilized by incubating in PBS supplemented with 1 % Triton (TX-100) overnight at 4°C and further blocked using 20% BSA for 4 hours. The permeabilized and blocked slices were then incubated by primary antibodies in 5% BSA-PBS incubated overnight at 4°C. After washing in PBS, slices were labelled with secondary antibodies conjugated with Alexa 405, 488, 555, or 568 for 3 hours at room temperature. Finally, slices were mounted on glass slides using DAPI fluoromount (Southern Biotech, Cat. No. 0100-20), as recently described^21^.

### Human Organotypic Slice Culture

Human neocortical slices were prepared as recently described^21,25^. Capillaries and damaged tissue were dissected away from the tissue block in the preparation medium containing Hibernate medium supplemented with 13 mM D+ Glucose and 30 mM NMDG. Coronal slices of 300 μm thickness were sectioned using a vibratome (VT1200, Leica Germany) and incubated in preparation medium for 10 minutes before plating to avoid any variability due to tissue trauma. Three to four slices were gathered per insert. The transfer of the slices was facilitated by a polished wide mouth glass pipette. Slices were maintained in growth medium containing Neurobasal (L-Glutamine) supplemented with 2% serum free B-27, 2% Anti-Anti, 10 mM D+ Glucose, 1 mM MgSO4, and 1 mM Glutamax at 5% CO2 and 37 °C. The entire medium was replaced with fresh culture medium 24 hours post plating, and every 48 hours thereafter.

### Chemical depletion of Microglia from slice cultures

Selective depletion of the myeloid cell compartment in human neocortical slices was performed by supplementing the growth medium with 11 μmol of Clodronate (Sigma, D4434) for 72h at 37ºC. Subsequently, the slices were carefully rinsed with growth medium to wash away any debris.

### Tumor/T cell injection onto tissue cultures

*ZsGreen* tagged BTSC#233 cell lines cultured and prepared as described in the cell culture section. Post trypsinization, a centrifugation step was performed, following which the cells were harvested and suspended in MEM media at 20,000 cells/μl. Cells were used immediately for injection onto tissue slices. A 10 μL Hamilton syringe was used to manually inject 1 μL into the white matter portion of the slice culture. Slices with injected cells were incubated at 37°C, 5% CO2 for 7 days and fresh culture medium was added every 2 days. Blood samples from the same donors from whom we obtained the healthy cortex for our organotypic slice cultures was drawn into an EDTA-cannula. Peripheral T cells were isolated using the same MACSxpress® Whole Blood Pan T Cell Isolation Kit (Miltenyi Biotech) and erythrocytes were eliminated from the suspension using ACK-lysis buffer (Thermo Fisher Scientific). T cells were tagged using the Cell Trace Far Red dye (ThermoFisher Scientific) prior to injection into the slices. To block endogenous IL-10 receptor, the neutralizing antibody anti-IL10 hAB (R&D systems) were added to the cells at the concentration of 5 μg/ml.

### Enzyme linked Immunosorbent Assay

An enzyme linked immunosorbent assay (ELISA) was performed in order to measure cytokine concentrations of IL-2, IL-10, IL-13 and IFN-gamma in the cell culture medium 48h after T cell injection. The Multi-Analyte ELISArray Kit (Qiagen, Venlo, Netherlands; MEH-003A) was used according to the manufacturer’s instructions. Absorbance was measured using the Tecan Infinite® 200 (Tecan, Männedorf, Switzerland).

### Treatment of patient with JAK-inhibitor

A patient with a recurrent glioblastoma was treated with a daily dose of 40mg Ruxolitinib for 4 weeks. Before treatment, we confirmed the progress by a biopsy. The treatment was performed as a neoadjuvant therapy. After 4 weeks, the patient underwent a gross-total surgery and adjuvant Temozolomide therapy.

## Acknowledgement

DHH is funded by the Else-Kröner Stiftung and BMBF. We thank Manching Ku and Dietmar Pfeifer for their helpful advice. We acknowledge Biorender.com.

## Data availability

scRNA-Sequencing Data available: (in preparation), Accession codes: www.github.com/heilandd/. VisLabv1.5 https://github.com/heilandd/Vis_Lab1.5, NFCN Algorithm www.github.com/heilandd/NFCN, SPATA-Lab: www.github.com/heilandd/-SPATA-Lab. Further information and requests for resources, raw data and reagents should be directed and will be fulfilled by the Contact: D. H. Heiland.

## Supplementary Figures

**Supplementary Figure1**: a) Workflow and representative FACS images. b) UMAP representation of all detected cell types. c-e) Distribution of patients across all clusters and cell types. d) UMAP representation of all determined clusters (SNN) e) The graph representing the percentage of each patient in different clusters f) Signature genes of each cluster g-i) UMAP representation of Isolated T cells spitted into CD4+ and CD8+ cells and the total number of clusters (13). j) Heatmap of signature genes

**Supplementary Figure2:** a) Copy-number alterations based on single cell data. Only a small subset of tumor cells was found in the OPC cluster. b) Gene expression maps of common marker genes.

**Supplementary Figure3:** a) Workflow to build a library of stimulated T-cells b) T cell stimulation in order to build a library for cytokine effects, illustrated is a heatmap of the 10 most significant marker genes of each stimulation state, based on PAMR algorithm implemented in the AutoPipe. c) Dimensional reduction (UMAP) of gene expression of the different simulation experiments. d) Z-scored expression of each stimulation signature along the velocity trajectory 1 (Figure 1).

**Supplementary Figure 4:** a) Workflow and concept of the NFCN Analysis

**Supplementary Figure 5:** a) Results from the Nichnet algorithm, left: The receptor-target interaction, right: the receptor-ligand network.

**Supplementary Figure 6**: a) Kaplan-Meier survival estimation of HMOX1 high/low expression GBM. b-Expression of HMOX1 in different regions of the tumor (b) and in de-novo and recurrent stage (c).

**Supplementary Figure 7**: a) Illustration of the workflow to determine the spatial correlation using a deep autoencoder for denoising followed by a Bayesian correlation model. b) Examples of the predicted overlap of T cells and CD163/HMOX1(+) myeloid cells. C) Distribution of the predicted correlation.

