## Supplementary figures and images for "Lineage and Spatial Mapping of Glioblastoma-associated Immunity"

### Supplementary Fig 1

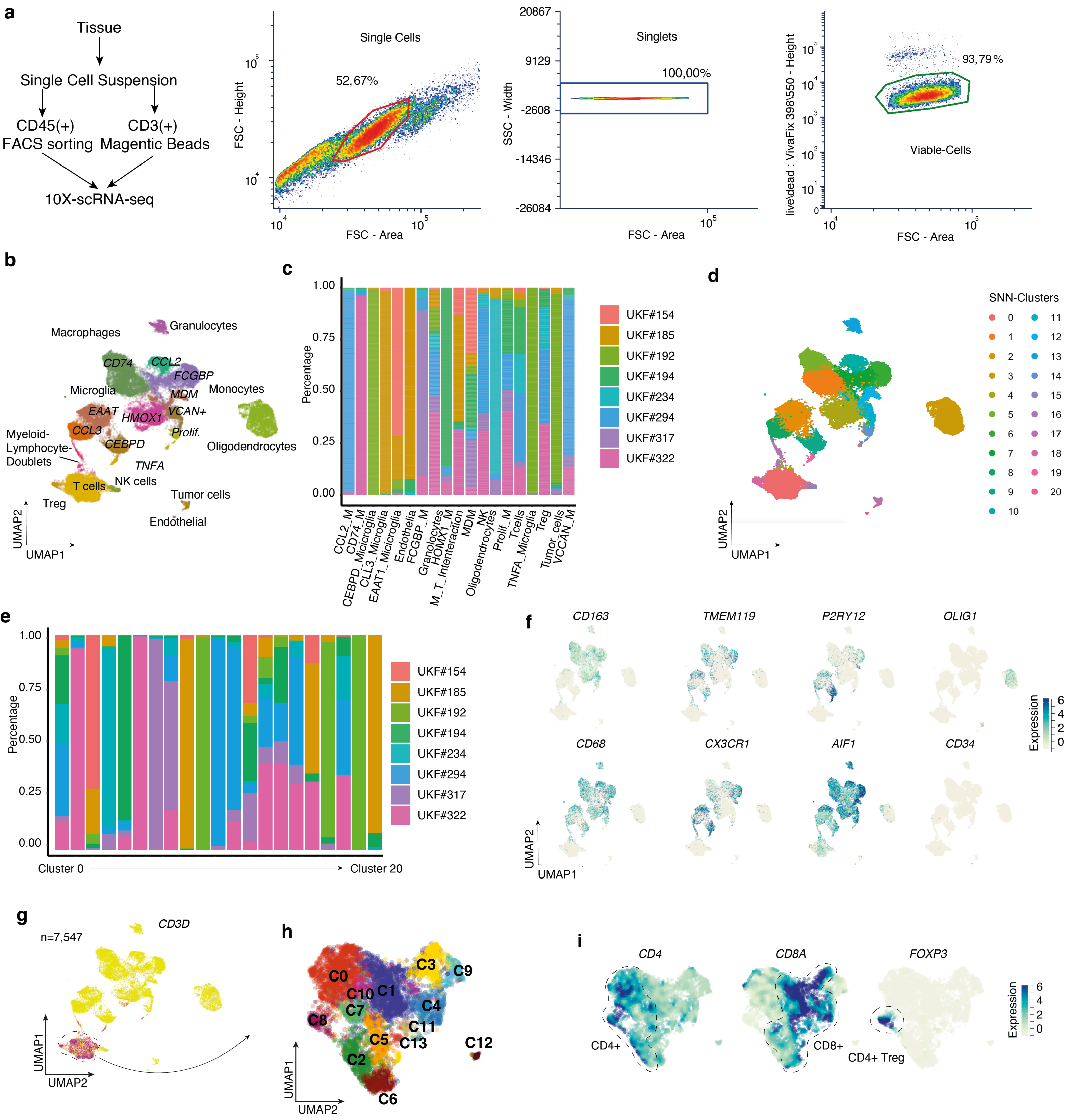

### Supplementary Fig 2

#### Supplementary Figure 2

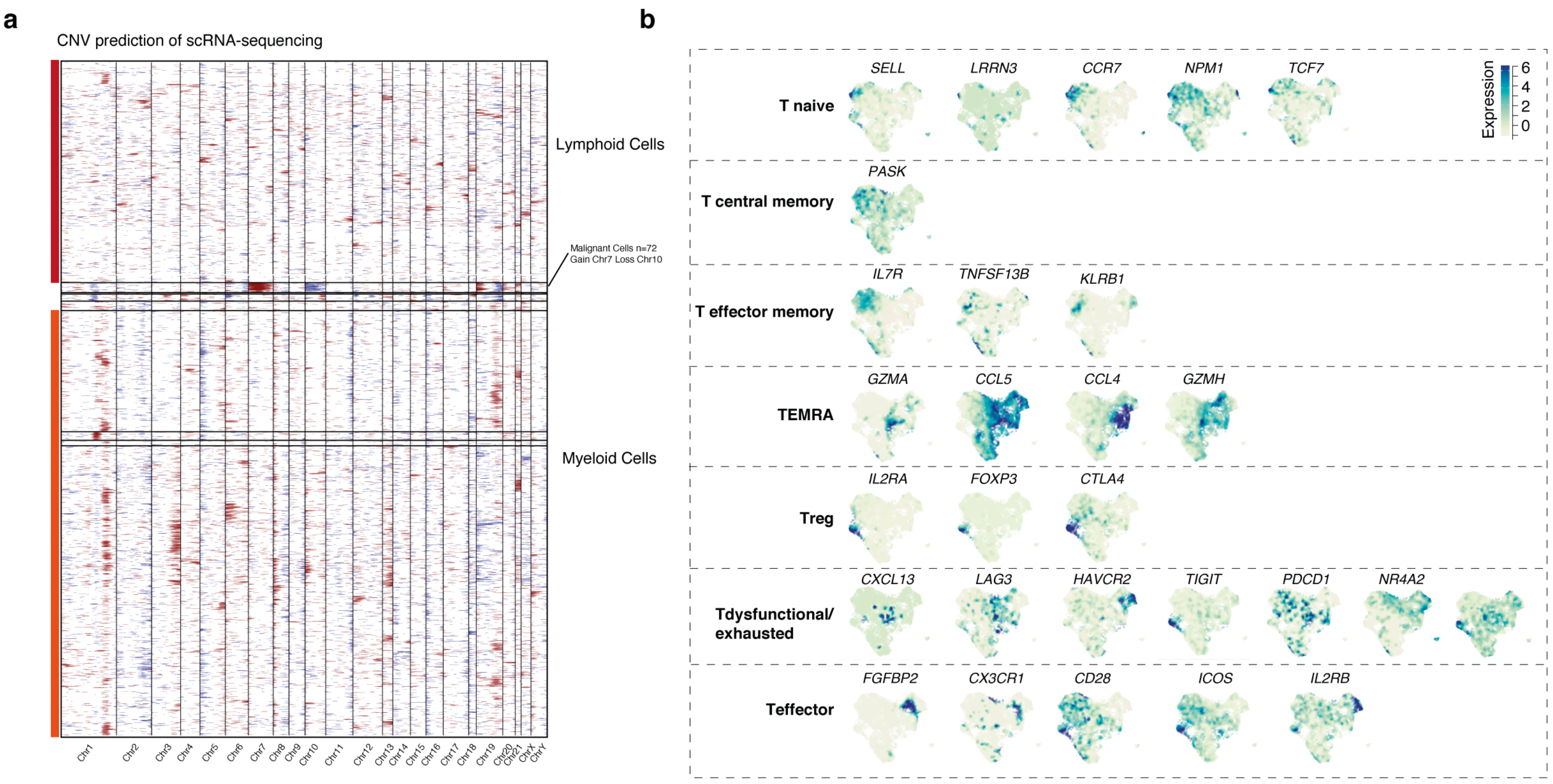

### Supplementary Fig 3

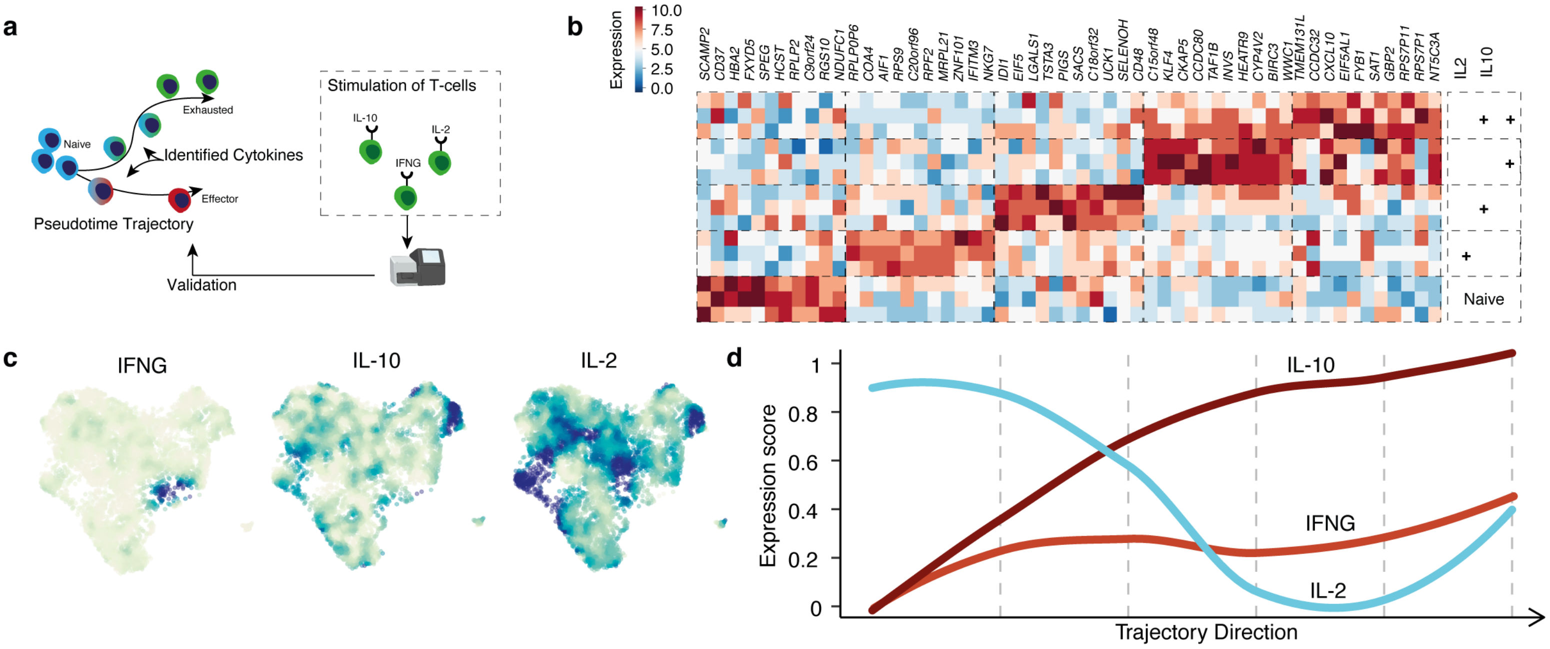

### Supplementary Fig 4

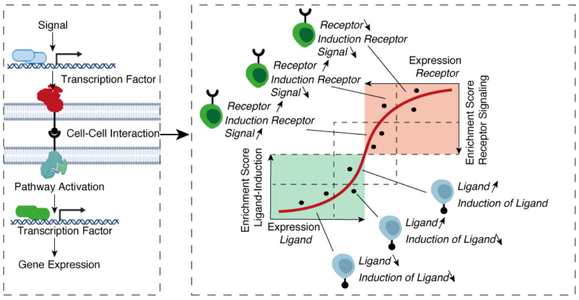

### Supplementary Fig 5

##### Nichnet Prediction of R-L Interaction

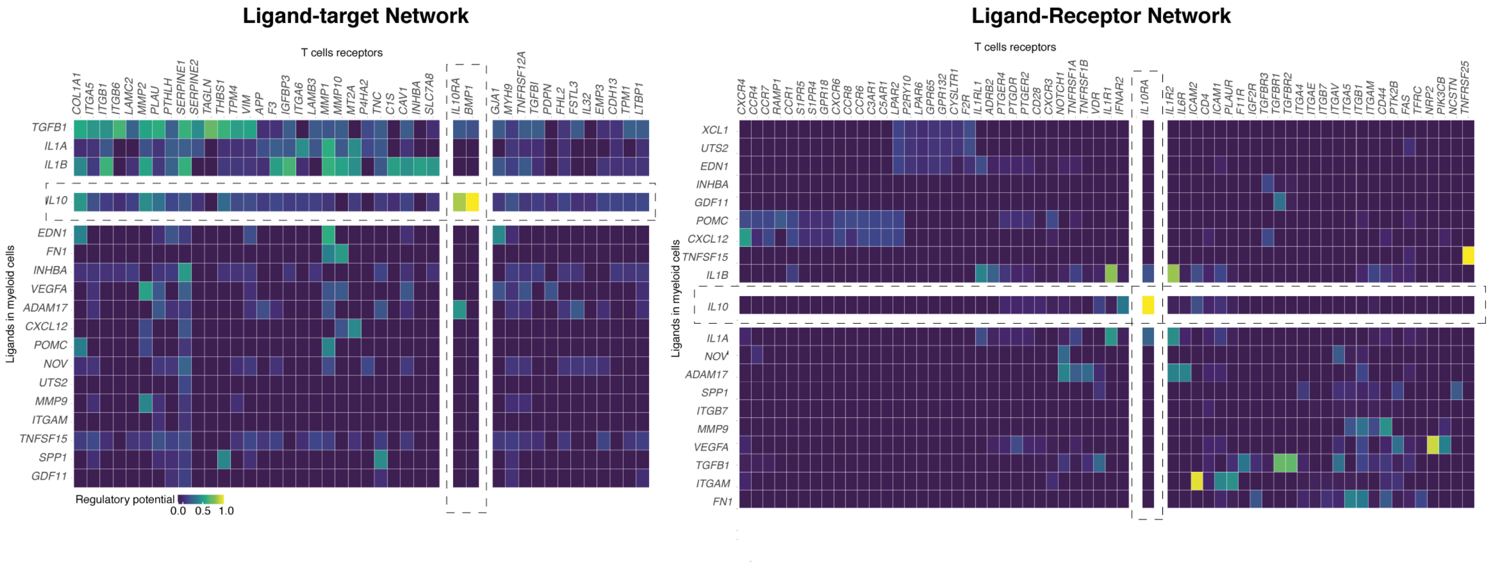

### Supplementary Fig 6

#### Supplementary Figure 5

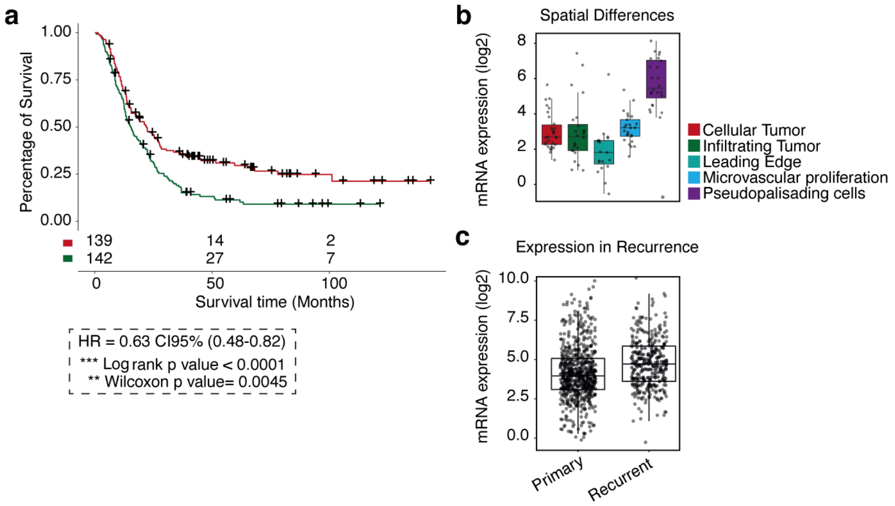

### Supplementary Fig 7

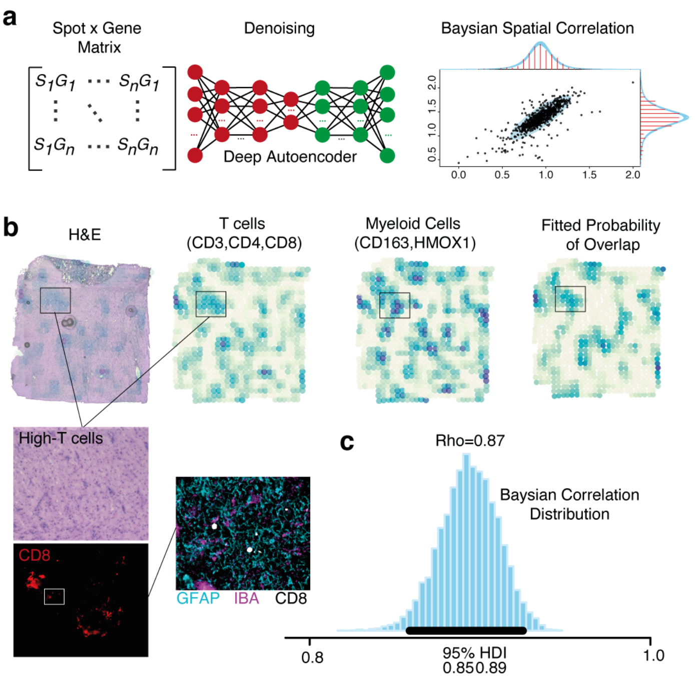
